# A model of bidirectional interactions between complementary learning systems for memory consolidation of sequential experiences

**DOI:** 10.1101/2019.12.19.882035

**Authors:** Michael D. Howard, Steven W. Skorheim, Praveen K. Pilly

## Abstract

The standard theory of memory consolidation posits a dual-store memory system: a fast-learning fast-decaying hippocampus that transfers memories to slow-learning long-term cortical storage. Hippocampal lesions interrupt this transfer, so recent memories are more likely to be lost than more remote memories. Existing models of memory consolidation that simulate this temporally graded retrograde amnesia operate only on static patterns or unitary variables as memories. However, the mechanisms underlying the consolidation of episodes, which are sequential in nature and comprise multiple events, are not well understood. There is currently no computational theory for how memory replays of sequential experiences emerge dynamically during offline periods and how they are coordinated between the hippocampus and cortex to facilitate the stabilization of episodic memories. Further, there is evidence for a bi-directional interaction between the two memory systems during offline periods (Ji & Wilson, 2007), whereby the reactivation of waking neural patterns originating in the cortex triggers time-compressed sequential replays in the hippocampus, which in turn drive the consolidation of the pertinent sequence in the cortex. We have developed a computational model of memory encoding, consolidation, and recall for storing temporal sequences that explores the dynamics of this bi-directional interaction and time-compressed replays in four simulation experiments, providing novel insights into whether hippocampal learning needs to be suppressed for stable memory consolidation and into how new and old memories compete for limited replay opportunities during offline periods. The salience of experienced events, based on factors such as recency and frequency of use, is shown to have considerable impact on memory consolidation because it biases the relative probability that a particular event will be cued in the cortex during offline periods. In the presence of hippocampal learning during sleep, our model predicts that the fast-forgetting hippocampus can continually refresh the memory traces of a given episodic sequence if there are no competing experiences to be replayed.

## Introduction

Studies on patients with lesions of the hippocampus report deficits both in forming new memories and in retrieving recent memories^1^. Based on these and subsequent studies, the hippocampus was found to be necessary for these functions^2, 3^. It learns rapidly due to its ability to quickly develop synaptic connections^4^, but fast learning also means fast decay as older memories are overwritten by new ones – which would mean catastrophic interference in a unitary memory^5–7^. This motivates the need for a two-stage memory system: the hippocampal engram gets strong quickly but also decays quickly, and the task of systems-level consolidation is to gradually train the more stable, slower learning neocortex^8–18^.

Replays of neural activity during sleep, which were first directly observed in the rat hippocampus^19–21^, have been hypothesized to play a key role in memory consolidation^15, 16, 18^. During slow-wave sleep (SWS), the cortex exhibits synchronized slow-wave oscillations (SWOs) that are characterized by a rhythmic alternation between depolarized (“UP states”) and hyperpolarized states (“DOWN states”)^22^. Replays generally occur during SWS^23^ and are coordinated between the hippocampal and cortical UP states^24^. Hippocampal replays can boost the weaker cortical representations and strengthen them until, eventually, they become independent of the hippocampus. Moreover, replays occur in a time-compressed manner^23–25^ to likely facilitate more efficacious strengthening of synaptic connections among pertinent neurons in the cortex for memory consolidation^23^ through spike timing dependent plasticity (STDP)^26–28^. The likelihood that a particular memory is replayed during limited SWS UP states appears to be dependent on saliency factors such as novelty^29^, recency, emotional involvement, and reward^30, 31^.

In the last 15 years, several animal and human studies have demonstrated improved post-nap or post-sleep memory performance by either boosting cortical SWOs with sensory or transcranial electrical stimulation^32–34^, or by applying cues during SWOs that were previously used to tag associations during learning^35–38^. These demonstrations suggest that even though cortex learns slowly, it can nonetheless be modulated to boost the memory consolidation process at various levels of specificity. Some experimental evidence suggests that neural activity in the cortex during SWO UP states leads and can potentially bias the content of the time-compressed replays in the hippocampus, which then in turn drive coordinated replays in the cortex^24^.

Existing computational models of memory consolidation^15, 16, 18^ primarily simulate the behavioral effects of lesioning the hippocampus at different times after learning. But they use only static patterns or unitary variables to represent memories, though the episodes that we experience on a daily basis are sequential in nature and have a temporal aspect to them. Moreover, the effect of hippocampal learning during sleep replays on the persistence of memory traces in the supposedly short-term storage in the hippocampus has not been examined. In this regard, none of the existing models have investigated any scheme to prioritize the reactivation of individual events during the limited offline periods. And as such, there is no computational theory for how memory replays of sequential experiences emerge dynamically during offline periods and how they are coordinated between the hippocampus and neocortex to facilitate the stabilization of episodic memories^24^.

In order to overcome these limitations, we developed a model of memory encoding, consolidation, and recall for storing temporal sequences that explores the waking and sleep dynamics of the bi-directional interactions between the fast-learning hippocampus and the slow-learning cortex. Details of the model are in the Materials and Methods section. Our model simulates episodic memories as a temporal sequence of activations of items in both regions^39, 40^, where each item represents the conjunctive activation of a pool of neurons sensitive to the salient features of a particular event in the experience^41, 42^. The Experimental Results section provides results from four simulation experiments. First, we demonstrate how a 5-item temporal sequence can be initially encoded in the hippocampus and subsequently consolidated in the cortex. We show how items can be probabilistically reactivated in the cortex during simulated UP states of SWS based on their recency and frequency of use (i.e., salience metric), how fast replays in the hippocampus can be triggered by these item reactivations in the cortex, and how they cascade into coordinated replays in the cortex that drive efficacious learning for long-term storage. Second, we replicate the phenomenon of retrograde amnesia with lesions of feedback projections from the hippocampus to the cortex. Third, we show how a more salient 5-item sequence is preferentially replayed and consolidated compared to another 5-item sequence. Fourth, we investigate if hippocampal learning during offline replays is critical for episodic memory consolidation. Finally, the Discussion section compares and contrasts our model with previous models of memory consolidation and also makes several testable predictions.

## Materials and Methods

### Model of Memory Encoding, Consolidation, and Recall

Our core model consists of a hippocampal module and a cortical module (Figure 1), which are both simulated as recurrent networks capable of learning sequences of items. Items, which are constituents of various sequential experiences, are represented by individual neurons in either module. There is all-to-all excitatory and inhibitory connectivity within each module. In particular, each neuron receives fixed inhibitory projections from all the other neurons, and each neuron can learn directional excitatory projections to other neurons in response to the sequential presentation of items from the environment. As proposed by the complementary learning systems theory^15^, intra-hippocampal excitatory projections have a much higher rate of learning compared to the intra-cortical excitatory projections. In other words, the cortex is slow to encode sequences of items compared to the hippocampus. Further, the memory traces (namely, weights of excitatory projections) in the hippocampus decay to their baseline levels (zero) at a much faster rate in the hippocampus compared to the cortex. Neurons also have self-excitation as well as delayed self-inhibition to regulate activations. Further, there are item-specific bi-directional fixed excitatory projections between the hippocampus and the cortex. Each waking sensory experience is represented as a sequence of items in an input register, providing bursts of excitation that coincide with their presentation to the corresponding neurons in the cortex. It is the spread of activation through the learned excitatory projections that is responsible for recall of a previously experienced sequence. In the remainder of this paper, for simplicity, we use ‘item’ to also refer to the cortical and hippocampus neurons, which each represent a particular item.

**Figure 1.**
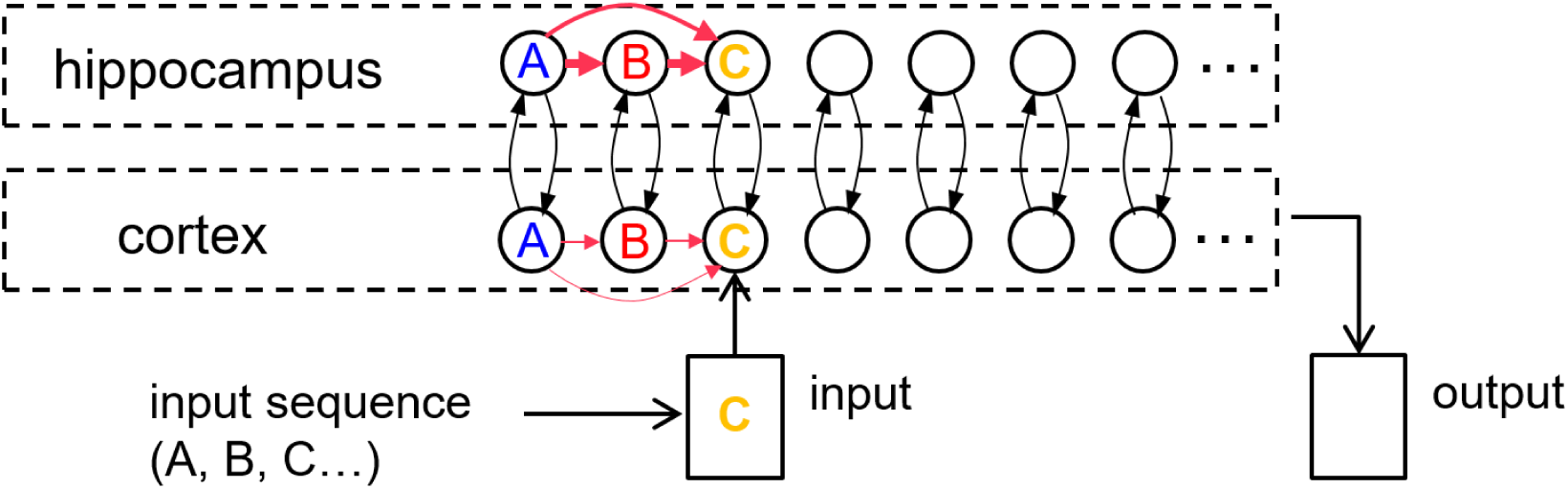
An illustration of the cortico-hippocampal memory model. Encoding, consolidation, and recall of sequential experiences are simulated. When a percept is entered into the input register, it energizes a matching cortical item. If a matching cortical item is not found, a new one is allocated. Activation is spread from cortical items to their counterparts in hippocampus and back to the cortex through fixed inter-module links or bi-directional projections (black arrows). Directional intra-module links (red arrows) develop between any simultaneously active items within each memory modulate as a function of their dynamic activations. The illustration shows an input 3-item sequence (‘A’-’B’-’C’) with directional links having formed first from ‘A’ to ‘B’. When ‘C’ is encountered, if ‘A’ and ‘B’ are still active, links are instantiated from ‘A’ to ‘C’ and from ‘B’ to ‘C’. The learning rate in the hippocampus is an order of magnitude faster than in the cortex.

Figure 2(a) is a plot of a training session in which the percepts ‘A’, ‘B’, ‘C’, ‘D’, and ‘E’ (applied to the input register for 2 s each) sequentially activate the corresponding items in the cortex, and thereby in the hippocampus. Figure 2(b) shows a recall cued by applying the energy of the single item “A” to the input register for 1.5 s, which spreads to activate items ‘B’ and ‘C’ in turn. Item activations in either memory module are ephemeral, falling soon after they rise to a high level. In the example shown, the network has not been trained enough to develop strong links to items ‘D’ and ‘E’ for a full recall. Figure 4 shows cued recall in more detail.

**Figure 2.**
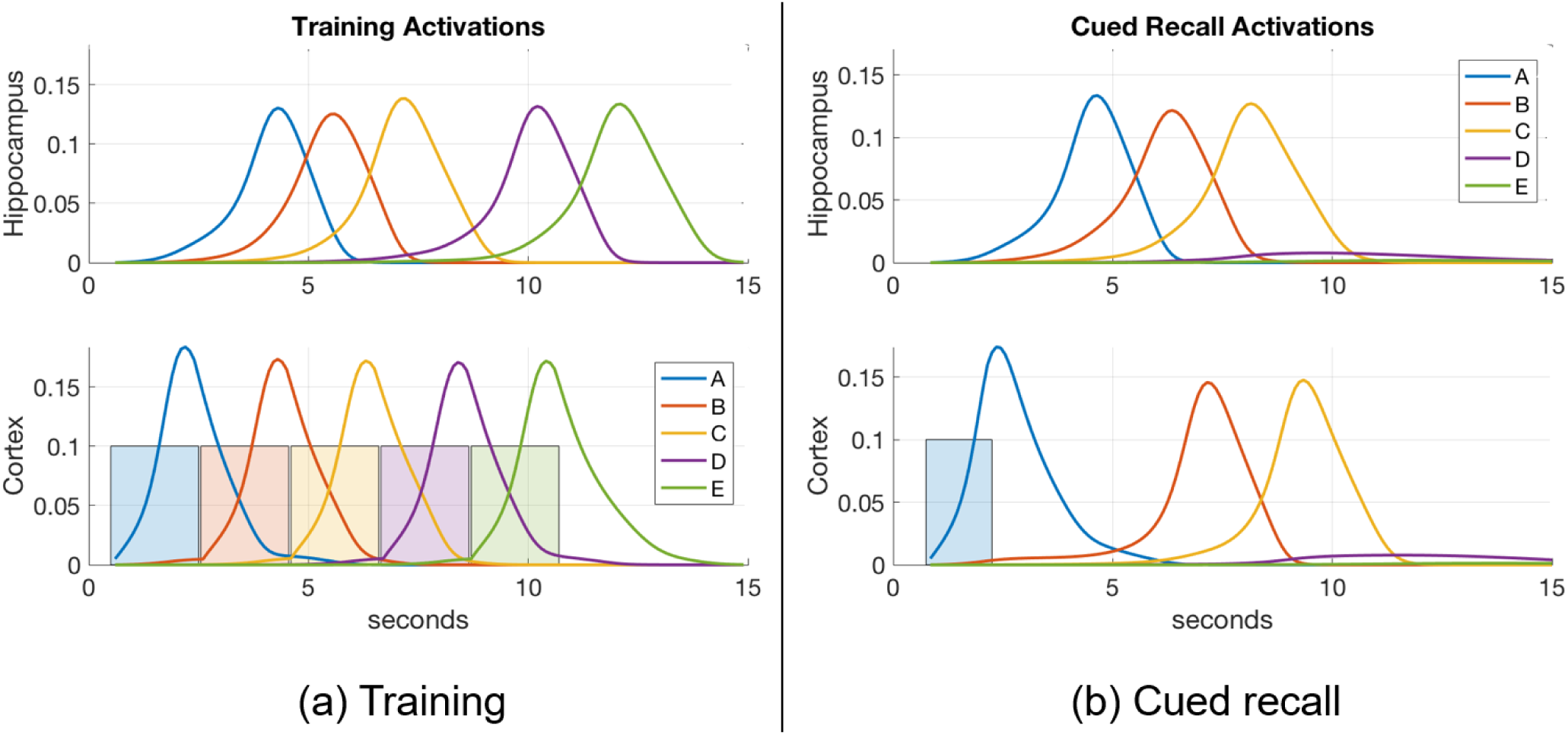
An example of item activations during (a) training and (b) cued recall. (a) A 5-item sequence is trained by presenting a series of inputs (the colored rectangles in cortex) that stimulate activation of matching items in cortex, and in turn activate the matching items in hippocampus. Directional excitatory projections between items within each memory module are learned as a function of their mutual activation dynamics. (b) Dynamics of neuronal activations when item “A” is cued in the input register after training. The activation of ‘A’ in hippocampus lags behind the cued activation in cortex as cortex is driving it, but subsequent items are activated more quickly in hippocampus due to its stronger excitatory projections (owing to faster learning). In the example shown, neither module’s weights have strengthened enough to spread activation through the entire sequence, so only ‘A’, ‘B’, and ‘C’ become activated.

**Figure 3.**
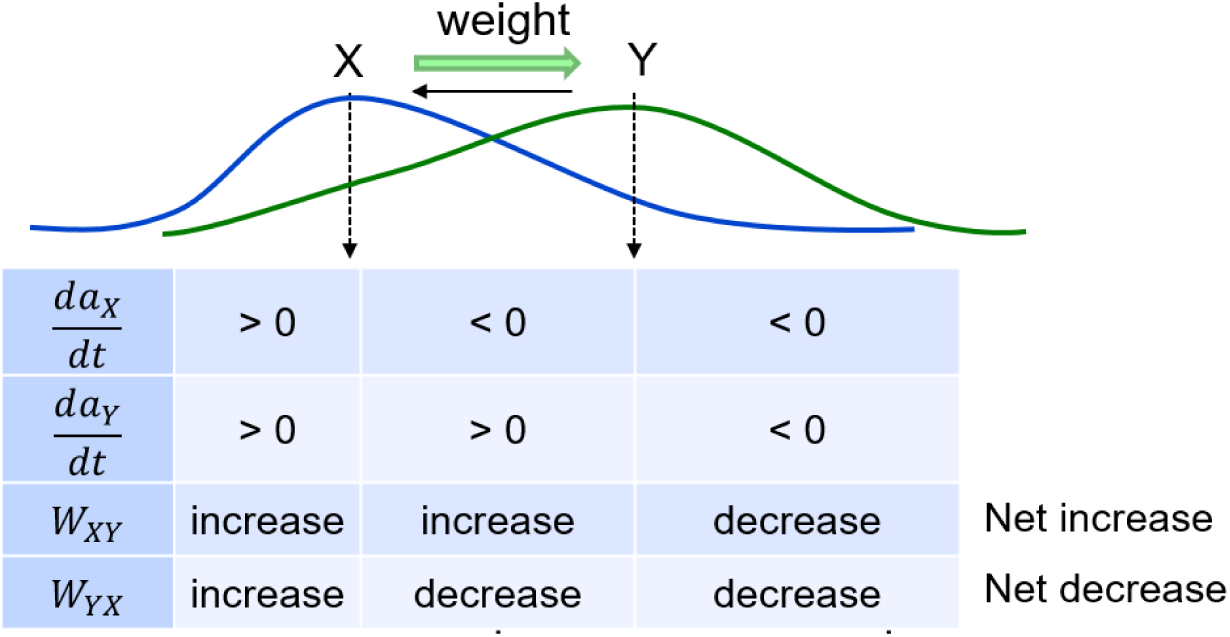
Weights develop as a function of item activations. Weights learn to be directional from earlier items to later items in a sequence.

**Figure 4.**
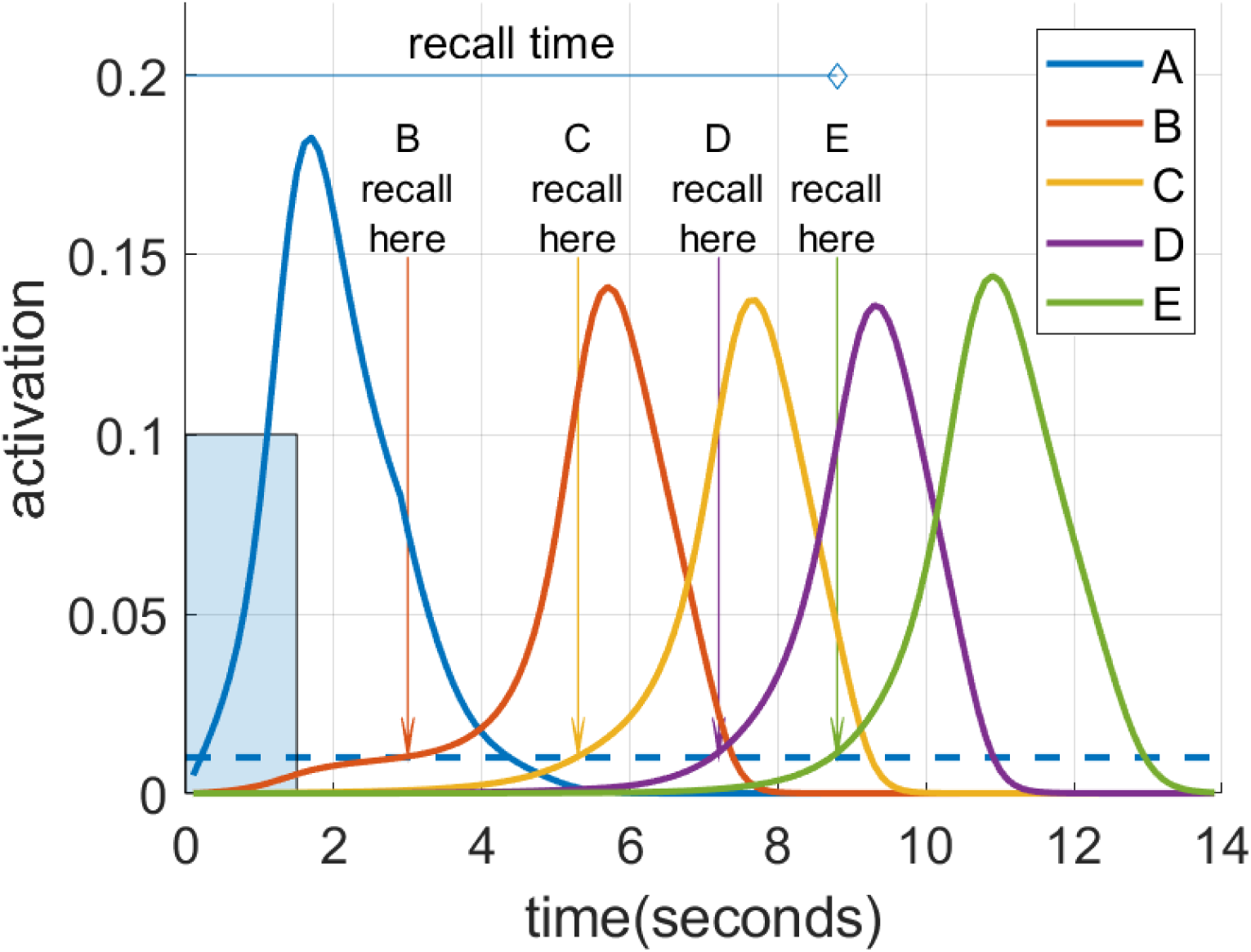
Spreading activation recalls a cued sequence during waking. After multiple training exposures, a presentation of the “A” cue (input activation of 0.1 for 1.5 s shown as a blue rectangle) causes a recall cascade in the cortex due to spreading activation through causal links. Each item in the sequence is considered recalled when it reaches the recall threshold of 0.01, shown as the blue dotted line.

### Simulation of memory encoding and recall during wake

The activation of a given item *x* in either cortex or hippocampus (*a*_*x*_), bounded between 0 and 1, is governed by shunting dynamics in a recurrent competitive network as follows:

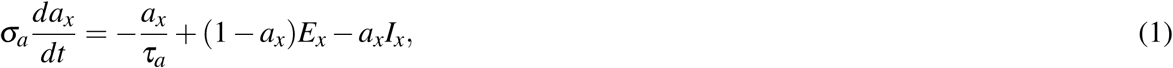

where *σ*_*a*_ modulates the rate of the neuron’s temporal integration (the smaller *σ*_*a*_ is, the faster the temporal dynamics are), *τ*_*a*_ modulates the rate of passive decay of the neuron’s activation (the smaller *τ*_*a*_ is, the faster the decay is), and *E*_*x*_ and *I*_*x*_ are the net excitatory and inhibitory inputs to the neuron, respectively. All model parameters are summarized in Table 1 along with default values. All equations were numerically integrated using Euler’s forward method with a fixed time step Δ*t* = 1 ms. The excitatory input to each neuron, *E*_*x*_, is a sum of the feedforward input *F* (from items in the input register to the cortex, and from items in the cortex to the hippocampus), the adaptive excitatory projections from other neurons in the module, and self-excitation (modeled using a sigmoidal function), as follows:

**Table 1.** Model parameters. Parameters with subscripts *h* and *c* correspond to different values for the hippocampus and cortex, respectively.

| Item activation<br>(Equations 1 - 3) | Description | Default Values |
| --- | --- | --- |
| $\sigma_a$ | parameter for speed of dynamics | $\sigma_{a,wake} = 2,$<br>$\sigma_{a,sleep} = 0.04$ |
| $\tau_a$ | passive decay parameter | 0.8 |
| $\mu$ | strength of feedforward excitation | $\mu_h = 2, \mu_c = 1$ |
| $\gamma$ | strength of spreading excitation between items | 0.9 |
| $\alpha$ | strength of self-excitation | 1 |
| $m$ | order of sigmoid nonlinearity for self-excitation | 2 |
| $t_a$ | threshold of self-excitation sigmoidal function | 0.09 |
| $\zeta$ | strength of feedback from hippocampus to cortex | 0.5 |
| $\beta$ | strength of inhibition between items | 15 |
| $\theta$ | strength of inactivation current | 10 |
| $m$ | order of sigmoid nonlinearity for inactivation current | 2 |
| $t_h$ | threshold of inactivation current sigmoidal function | 0.02 |
| Inactivation current<br>(Equation 4) | Description | Default Values |
| $\sigma_g$ | parameter for speed of dynamics | 10 |
| $\tau_g$ | passive decay parameter | 1.2 |
| $\kappa$ | strength of excitation from item activation | 1 |
| Adaptive weights<br>(Equation 5) | Description | Default Values |
| $\eta$ | learning rate | $\eta_h = 15, \eta_c = 1.5$ |
| $\tau_w$ | passive decay parameter | $\tau_{w,h} = 1.5552 \times 10^9,$<br>$\tau_{w,c} = 3.73248 \times 10^{10}$ |
| $Q$ | parameter for controlling non-causal links | 0.5 |
| $\eta$ | learning rate | $\eta_h = 15, \eta_c = 1.5$ |
| Item salience<br>(Equation 6) | Description | Default Values |
| $\tau_s$ | passive decay parameter | $\tau_s = 86400$ |
| $\lambda$ | gating variable for updating | $\lambda_{wake} = 1000,$<br>$\lambda_{sleep} = 0$ |

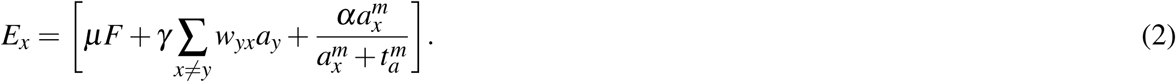

Note *F* = 0.1 for cortex (in response to the pertinent item in the input register) and *F* = *c*_*x*_ for hippocampus, where *c*_*x*_ is the activation of the corresponding item in the cortex. Cortical neurons are additionally excited by hippocampal feedback (*ζ* [*h*_*x*_]^+^, where *h*_*x*_ is the activation of the corresponding item in the hippocampus). The inhibitory input to each neuron, *I*_*x*_, is a sum of the fixed inhibitory projections from other neurons in the module and delayed self-inhibition (modeled using a sigmoidal function of an inactivation current *g*_*x*_), as follows:

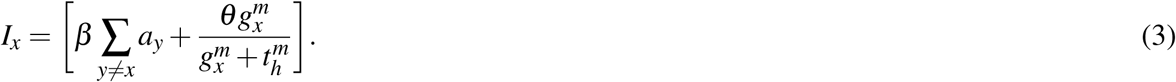

The inactivation current *g*_*x*_ simulates the inhibitory effect on active neurons that have been depolarized for a prolonged period^43, 44^ as follows:

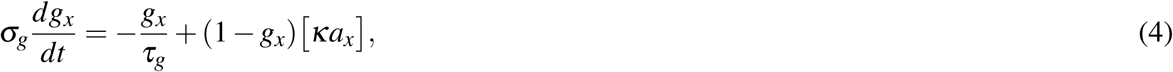

where *σ*_*g*_ modulates the speed of temporal dynamics and *τ*_*g*_ modulates the rate of passive decay. The inactivation current *g*_*x*_ evolves and decays at a slower rate than the corresponding neuron’s activation *a*_*x*_ (*σ*_*g*_ > *σ*_*a*_ and *τ*_*g*_ > *τ*_*a*_), providing delayed inhibition on the item when it gets large. The effect of self-excitation in Equation 2 is to raise the item’s activation quickly to a high level once activation reaches a certain threshold. However, the inhibitory input specified in Equation 3 ensures that few items will remain active at any time, and none for any prolonged period. All item activations *a*_*x*_ and inactivation currents *g*_*x*_ are initialized to 0 at the start of each experiment. Note that associative links between any two arbitrary items are instantiated at the first co-occurrence of their activations, as illustrated in Figure 1.

While an arbitrary item *y* is active, its energy spreads across weighted directional projections *w*_*yx*_ (in Equation 2) to other items *x*, and across the fixed links between the two memory modules (see Figure 2(b)). These directional projections are causal, based on the order in which the linked items are experienced. As a projection becomes stronger, more excitation is transmitted to the post-synaptic neuron, which becomes activated more quickly. This results behaviorally in faster memory recalls. Thus, the strength of the excitatory directional projections between the component items of a memory in both hippocampus and cortex are predictive of the ability to recall the sequential experience.

The adaptive weight of the excitatory directional projection from item *x* to another item *y* in either cortex or hippocampus (*w*_*x*_*y*), bounded between 0 and 1, is governed by the following rate-based learning rule:

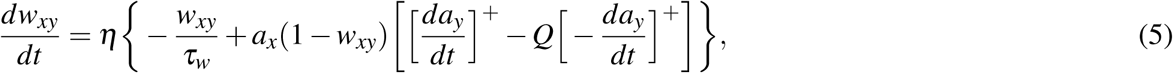

where *η* is the learning rate (the bigger *η* is, the faster the weights change), *τ*_*w*_ modulates the rate of passive decay, and *Q* is a parameter that controls the emergence of non-casual (reverse) links. Equation 5 models how causal associative links are formed as a result of overlapping activation of the pre-synaptic neuron (item *x*) and the rate of change in the activation of the post-synaptic neuron (item *y*) (cf., ^28, 45^). If the activation of item *x* reliably increases the activation of item *y*, then *x* is assumed to be causal and as a result the connection from *x* to *y* (namely, *w*_*x*_*y*) increases in strength. On the other hand, if the activation of item *y* reliably declines despite strong activation of item *x*, then *x* is assumed to be non-causal and as a result *w*_*x*_*y* reduces in strength. The reduction in weight is a fraction *Q* of the increase, which not only ensures that associative links in the causal direction are created, but also is necessary for the learning of connections between simultaneous events. At the default value for *Q* of 0.5, reverse links are eliminated. In the model, the hippocampus learns (*η*_*h*_ ≫ *η*_*c*_) as well as forgets (*τ*_*w*_, *h* ≪ *τ*_*w*_, *c*) faster compared to the cortex. With the default values for *τ*_*w*_ in Table 1, hippocampal weights undergo natural decay on the order of about 10 days and cortical weights on the order of about 2 years. Each weight is initialized to 0 when the corresponding connection is instantiated. Weights in both directions are updated at the same time, as illustrated in Figure 3. Existing connections are strengthened during recall, but new connections can only be created during training when a new item is presented while recently activated items are still active. Once associative links are learned between items in either memory module, a sequential memory can be reactivated by a relevant cue from the environment during waking or randomly during sleep. Note that by default, learning is enabled in both the hippocampus and cortex at all times including SWS.

### Memory recall metrics

Figure 4 illustrates the recall metrics used in the experiments discussed below, with an example of a cortical recall when cued by “A” after being exposed to the sequence ‘A’-’B’-’C’-’D’-’E’ (or ABCDE for short) for 20 trials. In this case, the cortical representation was strong enough that every item became activated in turn. An item is considered recalled when its activation level in the cortex rises above a threshold of 0.01. Recall accuracy is the proportion of sequence items successfully recalled in the correct order. So if all five items in the sequence reach the threshold in the correct order, the accuracy is 100%. However if any item crosses the threshold out of order, no credit is given for it or any further items. Thus if the activation level of the first and second items crossed the threshold in order, followed by the fourth then third and fifth, the accuracy would be 40% as two of the five items crossed the threshold in the correct order before an incorrect item recall. For a recall trial with 100% accuracy, the recall time is defined as the interval between the beginning of cue stimulation to the moment at which the final item crosses the threshold, provided this occurs before the arbitrary ceiling of 30 s. If the accuracy is less than 100%, the recall time is assigned the arbitrary ceiling value of 30 s.

### Simulation of memory consolidation during sleep

As described in the Introduction section, memories are consolidated in long-term cortical memory during SWS, when events experienced in the daytime can be reactivated or “replayed” during the UP states of SWOs. We simulate these emergent replays in each UP state by selecting an item probabilistically and activating it as described below. Figure 5 is a 6 s plot of emergent replays during a SWS simulation. An item cued in this way excites other items along existing directed connections, just as in waking, but the parameter for speed of dynamics (*σ*_*a*_ in Equation 1) is set so the activations rise and fall about 20 times as fast in sleep compared to waking. Sleep replays in rats are reported to occur between 6 and 20 times faster than waking experiences in rodents^23, 25, 46, 47^. There is also some evidence of time-compressed episodic memory replays in humans^48^. The actual speed of a given replay depends on the strength of the causal associative links between the constituent items of the corresponding sequence. The change in a given weight is primarily driven by the overlap in the activation of the pre-synaptic neuron and the rate of change in the activation of the post-synaptic neuron (see Equation 5), which is higher when the replay is compressed in time. Thus learning in the model is generally more efficacious during sleep replays compared to waking training. This is most important in the slower-learning cortex, and accounts for why sleep replays are so critical for strengthening connections in the cortex over time with no additional waking training.

**Figure 5.**
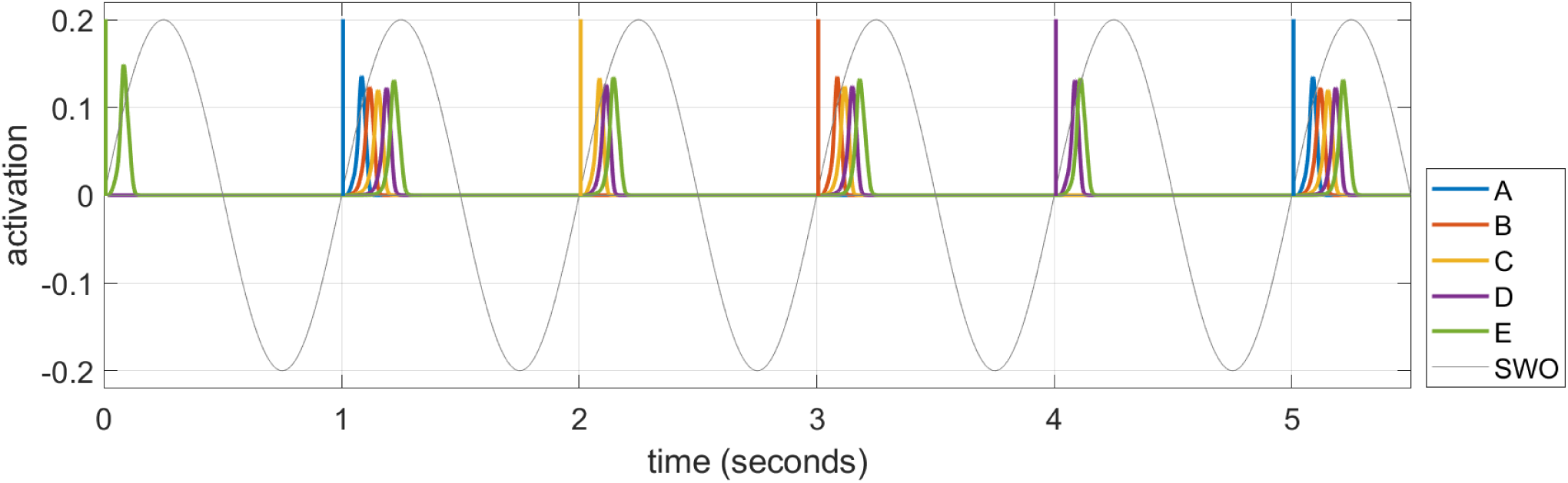
A series of time-compressed replays during UP states in SWS. The last 6 replays on Night 1 after training the sequence ABCDE for 10 trials are plotted. The initial item in each replay is selected from a weighted distribution based upon the recent activation of each item (i.e., the salience), and its energy spreads to the subsequent items of the pertinent sequence stored in memory. In this example, six replays are generated by cueing ‘B’, ‘C’, ‘C’, ‘E’, ‘D’, and ‘A’ in turn. The duration of the cue to initiate the replay in each UP state is 12.5 ms, and the amplitude is 0.2. A 1 Hz SWO is superimposed for illustration.

During UP states of SWOs, any item in either memory module can be randomly chosen to cue a replay by sampling from a weighted distribution called ‘salience’. The salience *s*_*x*_ of an item *x* is essentially a moving average of its activation during wake, which is defined as follows.

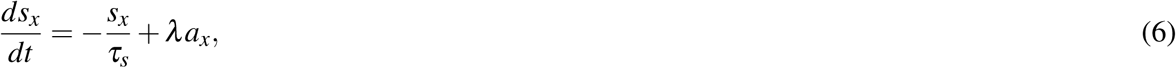

where *τ*_*s*_ modulates the rate of passive decay and *λ* scales the input of the corresponding activation, which is active only during wake. Activations during sleep replays do not contribute to salience in order to prevent a self-activation feedback that would lead to a few items dominating. Since any item can be selected, not just the first item in a sequence, replays are typically fragmentary. Note that all item activations *a*_*x*_ and inactivation currents *g*_*x*_ are reset to zero at the end of each UP state, which is also the beginning of each DOWN state.

## Results

### Experimental details

Figure 6 shows typical timings for presentation of input percepts and recall cues in the experiments that follow. Training trials present a sequence of input stimuli in the input register, such as the sequence ABCDE shown in Figure 2(a). Each percept during waking stimulates the input register at the level of 0.1 for 2 s, with an inter-stimulus interval of 5 ms. Trials are separated by 1 minute. When a recall test is required, it is separated from training by at least 1 minute, and the recall cue is presented in the input register for 1.5 s. As noted above, the recall is measured based on the spreading activations of the pertinent items in the cortex.

**Figure 6.**
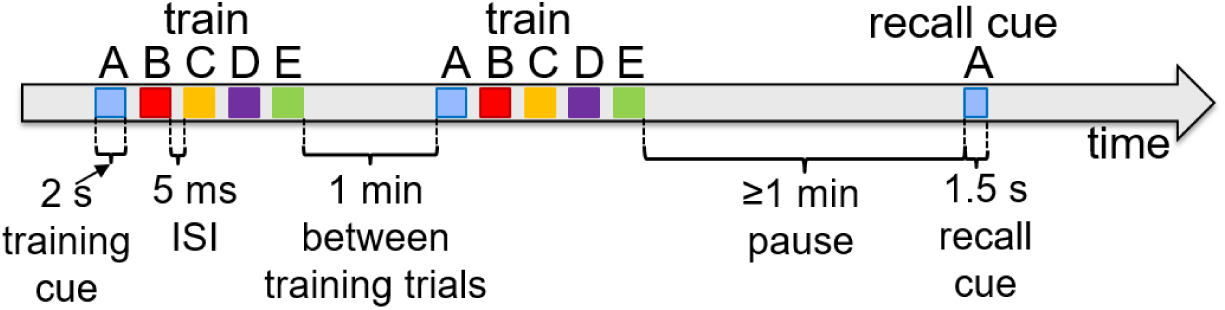
Typical experiment timings. Cue presentation times are illustrated for training and recall during waking. Training cues are presented for 2 s each in sequences, with 5 ms inter-stimulus interval. Recall cues last 1.5 s each.

For our simulation experiments, without loss of generality, each UP state during SWS randomly activates an item in the cortex with probability proportional to its corresponding salience with a brief pulsed cue in the input register (amplitude = 0.2, duration = 12.5 ms). In order to incorporate the possibility of no replay, a null item is added to the list of items with a fixed salience of 0.5. This ensures that the likelihood of selecting the null item is higher if the total salience of the other items is low (i.e., the probability of no replay increases when no items have been active recently). For simplicity and with loss of any generality, SWS during each night is simulated as a continuous block of SWOs for 50 s with a dominant frequency of 1 Hz. This manifests as 50 UP states per night, each with a duration of 0.5 s, to facilitate memory consolidation.

### Salience and memory replays

To show how different sequential experiences can interfere even when they do not share any items^49^ based on differences in practice during wake, we trained two different sequences (ABCDE and FGHIJ), 10 trials each on Day 1, and then only ABCDE for 10 trials per day on Days 2, 3, and 4. Each night, 50 replays were run during SWS using the salience algorithm described above to select from the 10 items (along with the null item) to cue for replay. Figure 7 shows simulations of the last full replay for both sequences in each night, which started with the pertinent first item in each sequence. The inset histograms show the number of replays cued by each item in the two sequences during each night.

**Figure 7.**
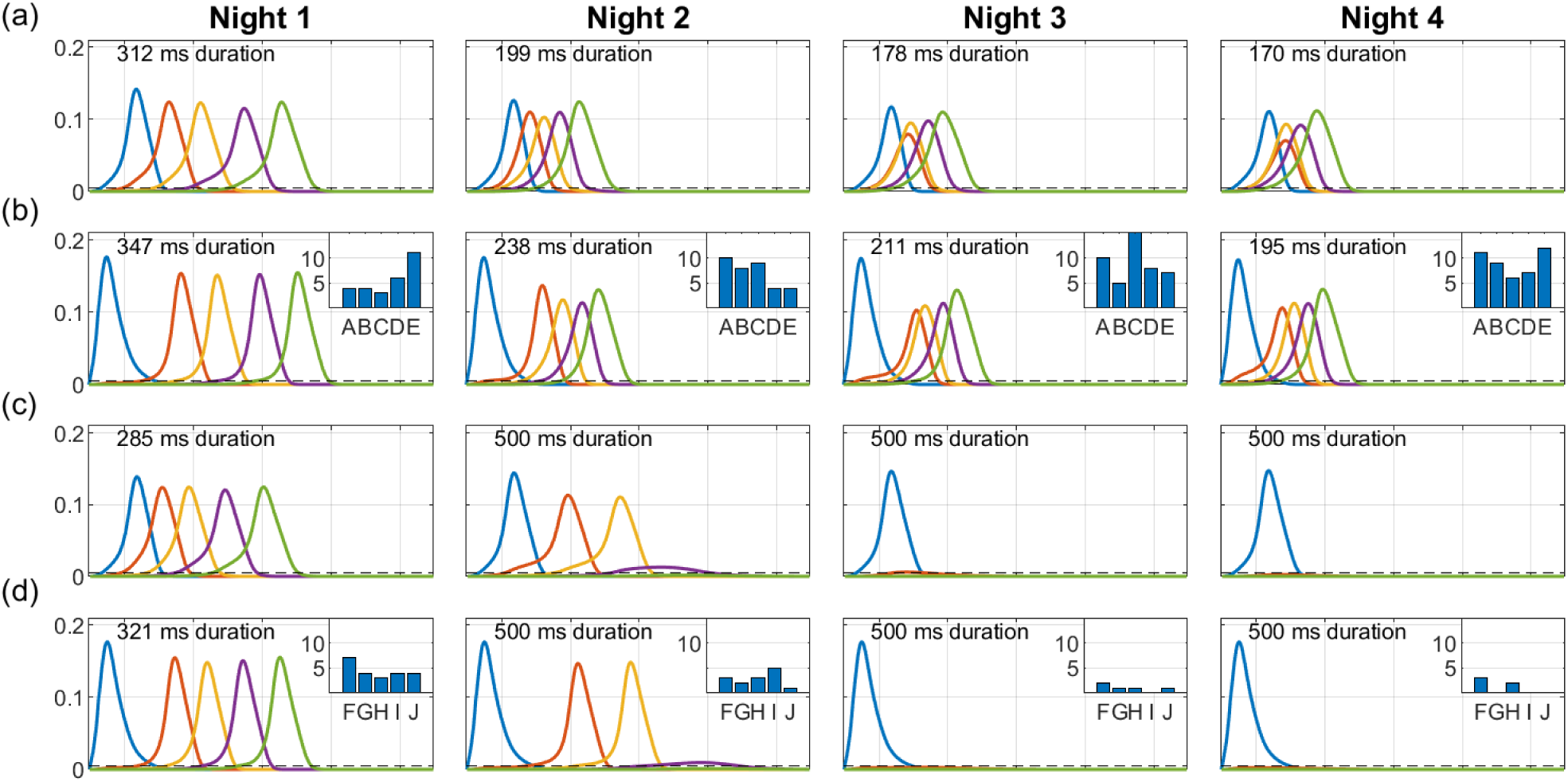
Representative replays of two separate sequences during a 4-day, 4-night simulation. Sequences ABCDE and FGHIJ were trained 10 trials each on Day 1, and then only ABCDE was trained for 10 trials per day on Days 2, 3, and 4. 50 cycles of 1 Hz SWOs were run each night. The last full replay of the night (cued by the first item in each sequence) is plotted here in context of the 500 ms UP state. (a) ABCDE replay in the hippocampus each night; (b) ABCDE replay in the cortex with inset histogram of the number of replays cued by each item during the night; (c) FGHIJ replay in the cortex each night; FGHIJ replay in the cortex with inset histogram of number of replays of each item during the night. Due to lower salience, replays of FGHIJ die out after Day 2 (the first item is activated by the replay cue). The duration of each replay is printed at the top of each plot. When all items in the sequence do not get activated, the duration is set to the length of the UP state.

Due to the extra daytime training on Day 2, the salience of the items in the sequence ABCDE rises above that for items of sequence FGHIJ, and so the sequence ABCDE is replayed more on subsequent nights (74% on Night 2, 94% on Night 3 and 98% on Night 4), and the activation spreads through the sequence faster with each night. Recall time of the replays is assessed at the end of the SWS period in each night by cueing with the first item for either sequence. The two sequences do not directly conflict in terms of sharing one or more constituent items, but they compete indirectly with each other for replay opportunities during SWS period in each night based on salience of the individual items. Additional training is certainly responsible for the increased number of replays for the sequence ABCDE on subsequent nights. Since there was no practice on the sequence FGHIJ after the first day, the salience of those items monotonically decayed; thus, the more salient ABCDE items were replayed more in the subsequent nights.

### Hippocampal influence on cortical replays

We use our model to explore the necessity of the hippocampus for the long-term consolidation of sequential experiences in the cortex using two simulation experiments. The first experiment reveals how the baseline network of intact hippocampus and cortex operates, and the second experiment focuses on what happens if the feedback projections from the hippocampus to the cortex are lesioned prior to the initial encoding of a sequence. For both experiments, the network was trained on the sequence ABCDE for 10 trials on Day 1, followed by memory assessment over the succeeding four days. This included four nights with a SWS period at the start of each night consisting of 50 UP state events where one item was chosen to cue the replay using the salience algorithm described above.

The dynamics of hippocampal and cortical weights are plotted in Figure 8 for the baseline case with both the hippocampus and cortex intact. Here the hippocampal weights grow to over 75% strength when trained on Day 1 and then quickly decay, whereas the cortical weights rise to less than 40%; however, the cortical weights suffer only imperceptible decay. At the beginning of each night, the hippocampal weights recover due to replays during SWOs, and the cortical weights rise as well. Replays occur in the hippocampus and the cortex on each night because the hippocampal weights are strong enough to propagate the activations through the sequence within the hippocampus, as well as within the cortex owing to item-to-item feedback projections from the hippocampus. During these replays, cortical connections become much stronger. Since there is no training after Day 1, the differences in the dynamics of weight strengths within each memory module over the subsequent days is exclusively due to the random emergent replays during the SWS period in each of the four nights; so, for example, in the cortex the weight *A* → *B* does not get as strong as the others because the fragment *A* → *B* within the sequence ABCDE is the least likely to be reactivated, compared to the other fragments, in response to saliency-based item selection for cueing replays during the SWS periods.

**Figure 8.**
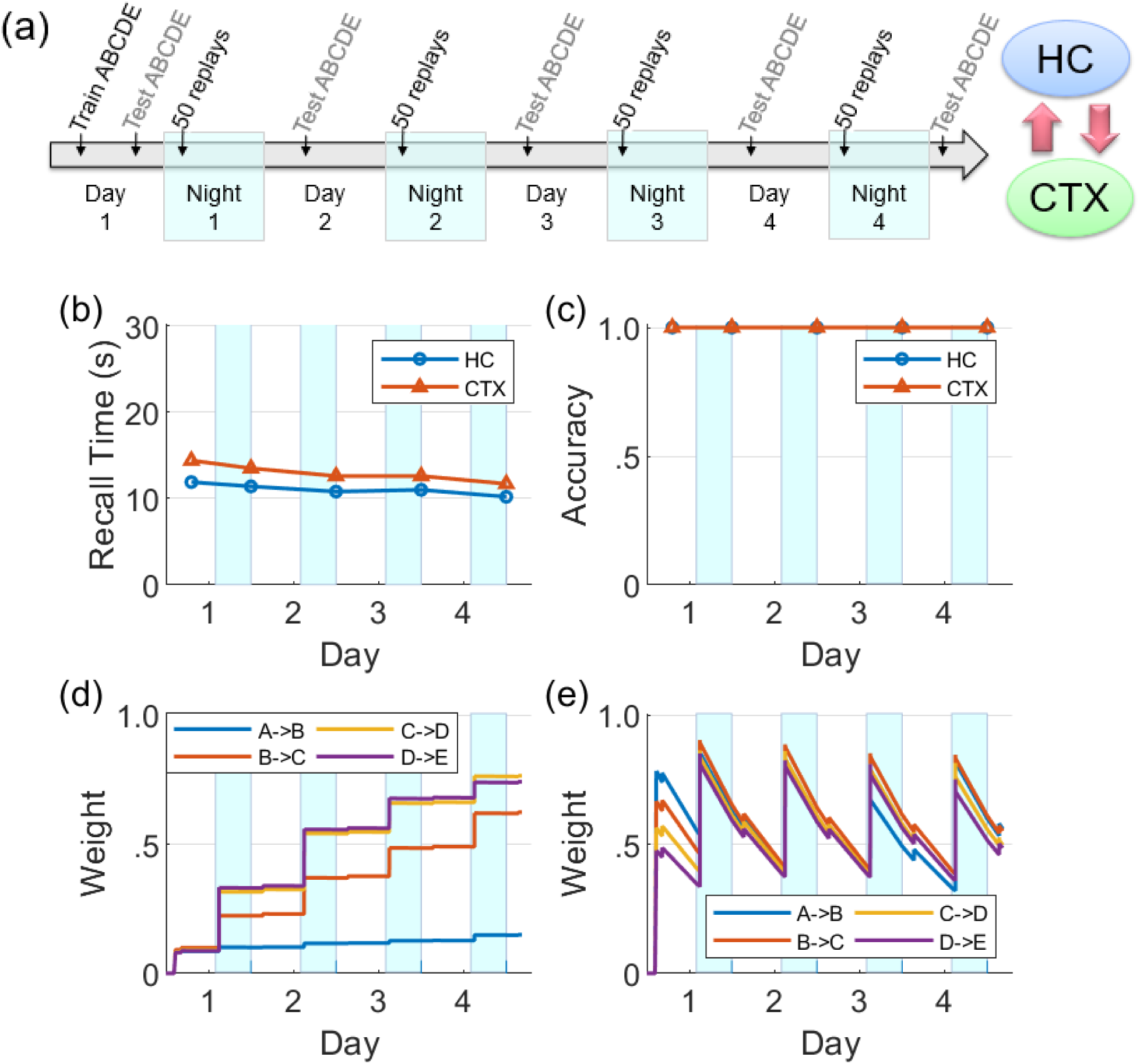
Baseline network. (a) A sequence ABCDE was trained and tested on Day 1 as illustrated in the timeline, and on the next four nights (highlighted with blue patches) there were 50 non-deterministic replays biased by salience, followed by morning tests (HC: hippocampus; CTX: cortex). The middle row plots the two memory recall metrics based on each of the five waking tests: (b) Recall time, where 30 s is default for incomplete defaults. (c) Recall accuracy. The bottom row plots the dynamics of weight strengths between items of the sequence ABCDE in the cortex and hippocampus during the same period: (d) Weight strengths in CTX. (e) Weight strengths in HC.

If the hippocampal projections to the cortex are lesioned prior to Day 1, we find that cortical replays cannot occur (Figure 9). Replays can still occur in hippocampus, but connections between the items in the cortex are not strong enough after the initial training to trigger replays on their own, and the cortical weights do not grow (cf., Figure 8). This is in agreement with the data on temporally graded retrograde amnesia with hippocampal lesions^14^, where the longer the hippocampus is intact after training (with intervening offline periods), the stronger is the long-term consolidation in the cortex. The hippocampal projections to the cortex are initially necessary to generate replays in the cortex during sleep, and thereby consolidate sequential memories in long-term storage.

**Figure 9.**
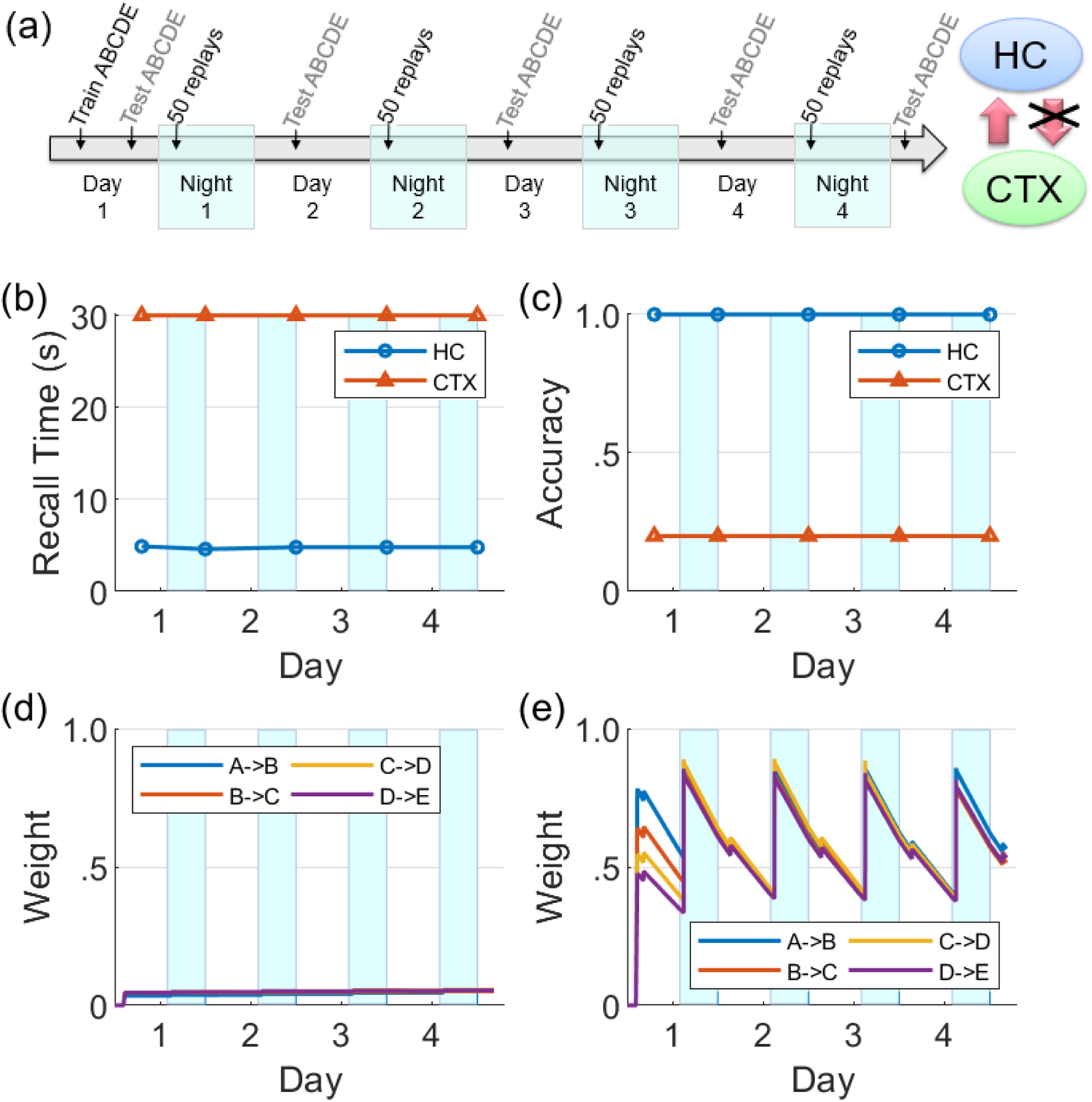
Hippocampal lesions before training. (a) With feedback from hippocampus to cortex turned off after training, the cortical representation cannot develop. Hippocampal connections are re-learned from replay activity during the SWS periods, but little growth is seen in the cortical connections (HC: hippocampus; CTX: cortex). (b) Recall time, where 30 s is default for incomplete defaults. (c) Recall accuracy. (d) Weight strengths in CTX. (e) Weight strengths in HC.

### Competing sequences

We now focus on the effects of competition among sequential experiences for limited replay opportunities during sleep on memory recall. In this experiment (see Figure 10(a)), two non-overlapping sequences ABCDE and FGHIJ were trained for 10 trials each on Day 1, followed by continued training for the sequence FGHIJ only on subsequent Days 2-4 (10 trials per day). This has the same design as the experiment reported in Figure 7. Memory recall for the sequence ABCDE was assessed on each of Days 1-5 to focus on the effect of the more practiced sequence on the less practiced sequence. Just as the simulation experiments reported above, there were 50 replay opportunities at the start of each of the four intervening nights. On Nights 2-4, the items belonging to the sequence FGHIJ have a much higher chance (compared to the items belonging to the sequence ABCDE) of getting selected to cue the replays during the SWS period, as they have higher salience due to the additional practice on Days 2-4.

**Figure 10.**
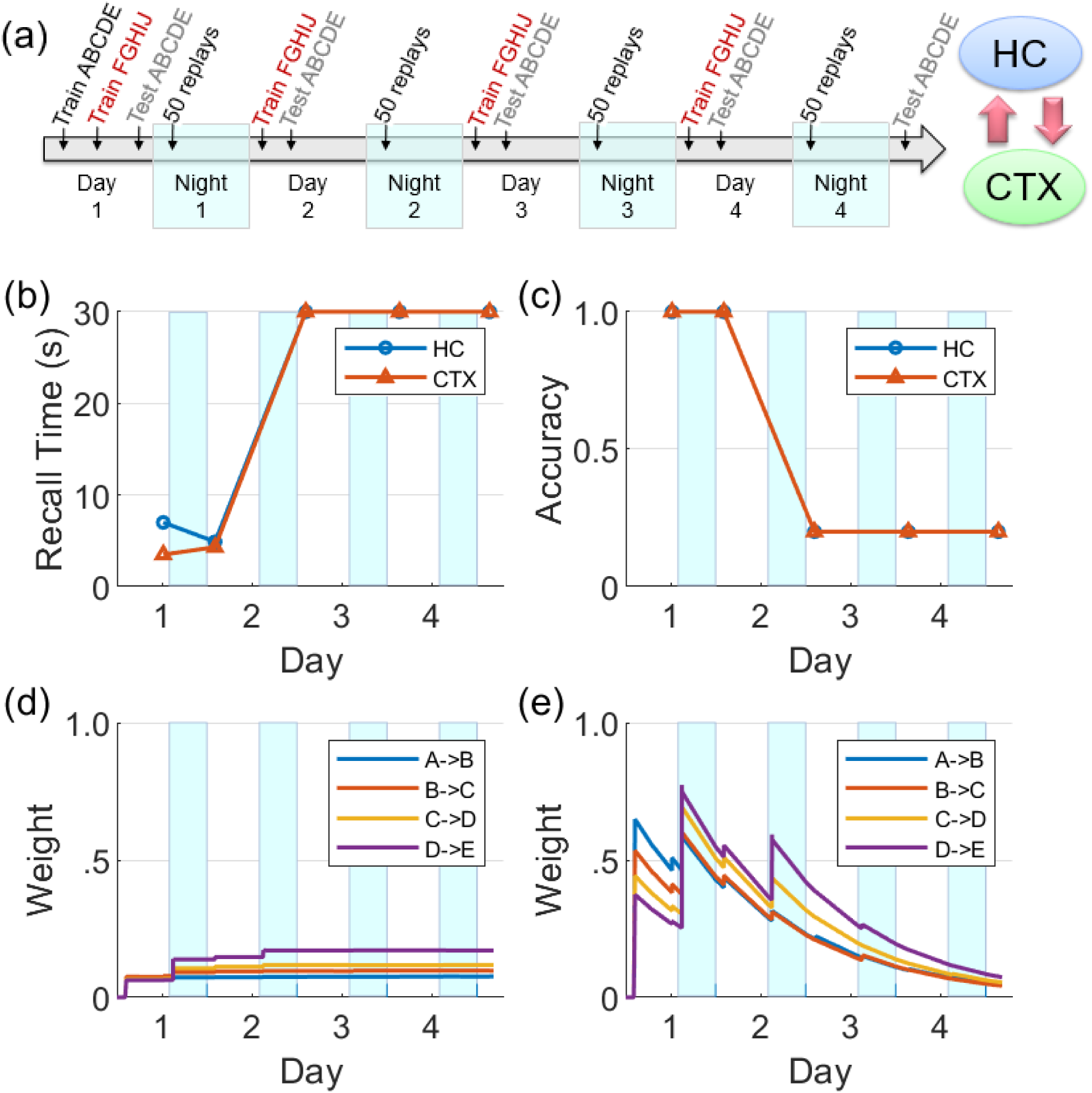
Effects of competition for limited replay opportunities on memory recall. This experiment is similar to Figure 8, but competition from the more practiced sequence FGHIJ reduces and then eliminates replays of the original sequence ABCDE (HC: hippocampus; CTX: cortex). There is a growth of cortical connections in Night 1, but very little in Night 2 and beyond. (b) Recall time for the sequence ABCDE, where 30 s is default for incomplete defaults. (c) Recall accuracy for the sequence ABCDE. (d) Weight strengths in CTX for the sequence ABCDE. (e) Weight strengths in HC for the sequence ABCDE.

In Figure 10, it can be seen that the cortical and hippocampal weights for the sequence ABCDE increased similar to Figure 8 on Day 1 and Night 1. But starting from Night 2, the preferential replays cued by items belong to the other sequence FGHIJ result in the indirect suppression of consolidation of the sequence ABCDE in the cortex. By Night 3, there were almost no replays related to the sequence ABCDE. Consequently, the sequence ABCDE is not recalled on Days 3-5 with expected minimum recall accuracy and maximum default recall time.

### Hippocampal learning disabled during sleep

There is evidence that hippocampal plasticity is suppressed during SWS^50, 51^. This might serve the purpose of allowing transfer to the cortex without further modifying the hippocampal representation^14^. We repeated the experiment reported in Figure 8 with hippocampal learning disabled during sleep. During Day 1 training, the hippocampus learned the sequence as evidenced by the high weights, high recall accuracy, and low recall time. Hippocampal replays occurred during Night 1, but without learning in sleep. Unlike Figure 8(e), the hippocampal weights decayed continuously following the initial encoding except for small bumps during daytime recall testing. However, in this case, the feedback from hippocampus to cortex was intact; so even as the hippocampal weights decayed without the benefit of the overnight replays, there was enough replay activation energy to strengthen the cortical weights during SWS to the point that the cortical representation became independent of the hippocampus. Compare with Figure 9 to see the difference between lesioning the hippocampal projections to the cortex versus having an intact hippocampus with learning disabled during sleep. This shows that while the hippocampal projections to the cortex are critical for long-term consolidation, learning in the hippocampus during SWS may not be critical.

## Discussion

We have presented a model of the cortico-hippocampal system that quickly learns causal associations between items in sequential experiences as a function of the overlap of their dynamically changing activations for the short term, and over a longer period of time develops a stable long-term representation. The model produces memory consolidation due to emergent replay activity during SWS periods^10, 12–14^ (see Figures 8 and 11). As illustrated in Figure 1, our model demonstrates the key role of the bi-directional cortico-hippocampal dialogue to strengthen and consolidate sequential memories for long-term storage in the cortex. Figure 8 shows that with hippocampal learning intact during sleep if the model doesn’t experience new memories, the hippocampal representation of a given sequence can sustain itself almost indefinitely (or at least until the salience of the items belonging to the sequence decays to an extremely low level). Figure 9 shows that if hippocampal projections to the cortex are lesioned, the cortex is unable to develop a long-term representation. For the case that the hippocampal projections to the cortex remain intact, if there is increased competition from another sequence (due to additional practice) for replay opportunities during SWS periods before the cortical representation is developed, the less practiced sequence is not replayed enough in the hippocampus to facilitate consolidation (see Figure 10). On the other hand, plasticity is suppressed in hippocampus during SWS^50, 51^, possibly to prevent changing the hippocampal representation while its replays are driving the transfer to the cortex. If the network remains intact but hippocampal learning is turned off during sleep, the cortical representation can develop even as the hippocampal representation decays (see Figure 11).

**Figure 11.**
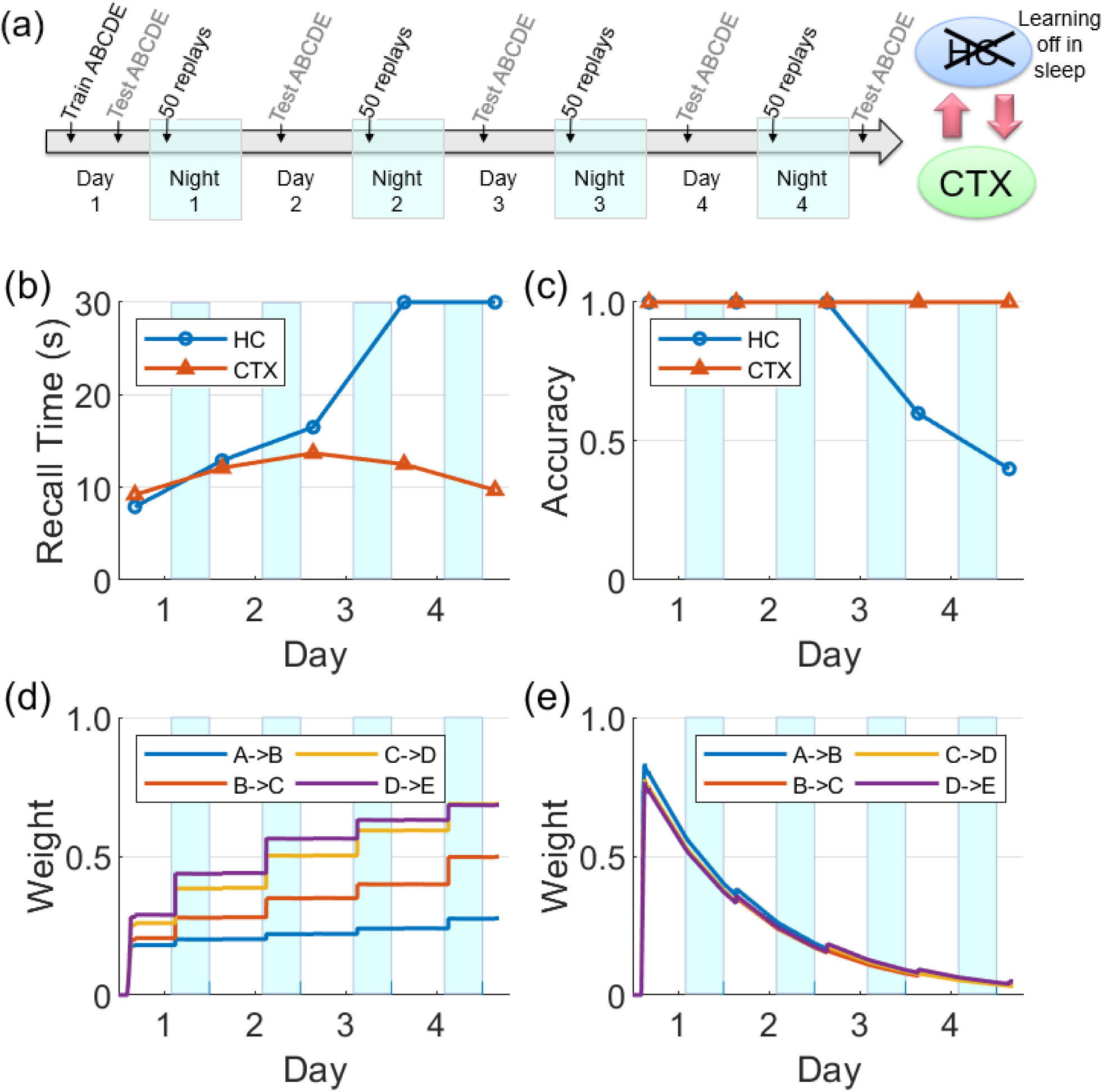
Intact network with hippocampal learning turned off during sleep. Hippocampal replays occur during sleep, but with learning turned off the hippocampal weights do not benefit and decay continuously. However, sufficient weight strengths are built up in the cortex during Night 1 such that the cortex is able to sustain replays through the subsequent nights. In this case, the cortex can still consolidate the sequence and become independent of the hippocampus.

### Replays and salience

Replays are primarily observed during the UP states (500 ms) of SWOs during SWS^23^, which we have simulated as shown in Figure 5. Replays have also been detected during quiet waking^52, 53^. However, it has been observed that replays during SWS are more effective for consolidation than waking replays^54^. This may be due to powerful bursts of activation during sharp-wave ripples in the hippocampus during SWS^55^, which speed up the dynamics of the sleep replays as much as 20 times over the original waking experiences^23–25, 46, 47^. In the brain, the temporal compression of replays means a tighter overlap between the activations of the items leading to more efficacious learning of the pertinent connections in the cortex. It also allows longer range connections, so that events that were too temporally separated during waking can be associated during sleep^56, 57^. Plasticity is suppressed in the hippocampus during SWS^50, 51^, but not in the cortex, so replays during sleep give the slower learning cortex more exposures to a memory at a higher effective learning rate than during the original experience. Note that in the experiment of Figure 11, even with hippocampal plasticity disabled during sleep, the memory can still be transferred to cortex successfully.

Do replays start at the beginning of a sequence, or more often in the middle due to random reactivation? Do they continue reliably to completion? Existing physiology data^19, 23, 24, 53, 58^ correlate neural spiking during replays with entire movement sequences and can tell whether a waking experience was replayed in the forward or reverse direction, but does not clarify whether partial sequences were replayed. Our model simulates replays by activating any item in memory with a probability proportional to its salience metric (see Equation 6). If items for a given sequence were randomly chosen to cue replays and the activation always spread in one direction to the end of the sequence, the weights of the later links would grow stronger than the earlier links since it is more likely they would be adapted over the course of many replays^59^.

Without the influence of salience, old and new memories would all replay with equal frequency, reducing the opportunities for a new memory to become consolidated. The salience metric defined in Equation 6 makes it more likely for newer memories to replay. In our model, the salience of a memory is only updated during waking experience; if salience were also updated during sleep replays, runaway feedback would prevent new memories from competing for consolidation. We know that salience is not a simple function of recency and can be influenced by neuromodulators^60^. It is possible that the likelihood of replays may be reduced due to physiological factors during encoding such as low attention or high fatigue, or increased by novelty^29^, high emotional content^61^, or immediate reward^30, 62^. In this regard, McNamara and colleagues^31^ showed that optogenetic stimulation of dopaminergic neurons in rat ventral tegmental area (VTA) during spatial learning improves offline hippocampal replay strength by approximately 40%, in terms of correlation between wake and sleep firing patterns of hippocampal cells coding novel spatial memories. Such neuromodulation makes it possible for statistically infrequent but task-relevant events to become preferentially consolidated in the cortex.

### Previous memory consolidation models

Several other computational models of memory consolidation have been published. But they use only static patterns or unitary variables to represent memories, do not consider the temporal aspects of sequential episodes, do not model the emergence of time-compressed replays that are coordinated between the hippocampus and the cortex during offline periods, and have not investigated the role of hippocampal learning and the effect of limited replay opportunities during offline periods. Alvarez and Squire^18^ simulated memory transfer from short-term to long-term storage using a cortical-hippocampal neural network model with inter-areal projections but without intra-areal projections. By lesioning the hippocampal layer after training, they produced temporally graded retrograde amnesia in which memories are impaired only to the extent that they had not been consolidated before the lesion. McClelland and colleagues^15^ simulated the phenomenon of memory consolidation using a short-term hippocampal storage and long-term cortical storage, which were implemented as variables of memory strength. As a result of memory encoding, the hippocampal memory strength is instantiated at a higher level than the cortical memory strength. Further, the memory strength in the hippocampus decays at a faster rate compared to that in the cortex. Cortical memory strength is boosted during offline periods by the phenomenon of reactivation in proportion to the corresponding memory strength in the hippocampus. Memory recall accuracy is modeled as a function of memory strengths in both the hippocampus and the cortex. Kali and Dayan^16^ simulated a multi-modal hierarchical associative network with reciprocal projections in the slow-learning cortex and a hippocampal buffer of novel patterns from the conjunctive layer in the cortex. During offline periods, the hippocampus reactivates its memories randomly in the cortical conjunctive layer to facilitate more learning and thereby consolidation in the cortex.

### Do hippocampal memory traces survive long term?

Figures 8 and 9 show a hippocampal representation that survives through the course of the experiment (four days) with no sustained decay. However, in those experiments there is no competition for limited replay opportunities during SWS periods from other sequences. And although the hippocampal memory traces decay rapidly each day (see Equation 5), they are refreshed by replays each night due to hippocampal learning that is enabled at all times. However, in the experiment where the hippocampal learning is disabled during SWS, the hippocampal representation does not survive for long (see Figure 11) because the weights are passively decaying at a high rate without any updates from the replays. Our model supports a systems-level consolidation theory in which a hippocampal representation is effectively transferred to the cortex^8–18^. A related suggestion^18, 63^ is that offline memory consolidation serves to bind together multi-modal aspects of an episode by developing cortical links between “cross-cortical storage” fragments so that a distinct episodic memory can be recalled.

It is generally accepted that cortical representations are more semantic, more generalized^64^. So it is not clear how certain detailed remote episodic memories can be recalled for many years if the hippocampal representation decays completely. Multiple trace theory (MTT)^65^ suggests that a hippocampal representation survives as long as the memory can be recalled, and is the source of detailed episodic memories that cannot be provided by the generalized cortical representations. The Kali and Dayan model^16^ supports that theory by demonstrating that if a hippocampal representation does persist, it is possible for it to stay linked to the corresponding cortical representation as it evolves over time. They speculate that the purpose of replays is to maintain the hippocampal indices by which the cortically stored memories can be retrieved. Nadel and colleagues^65^ also present fMRI evidence in support of MTT, which shows high hippocampal activation during recall for not only recent memories (which are dependent on the hippocampus) but also remote memories (which are presumably not dependent on the hippocampus). In our model, cueing the recall of a remote memory in the cortex would elicit the reactivation of the corresponding items of the sequence in the hippocampus due to projections from the cortex to the hippocampus even though the underlying weights have decayed significantly. In other words, just as hippocampal replays of unconsolidated, recent memories drive the replays in the cortex, the cortical recall of consolidated, remote memories drive the reactivations in the hippocampus. We suggest that the phenomenon of reconsolidation^66, 67^, wherein so-called consolidated memories become temporarily labile following a recall and subject to modification, is facilitated by the hippocampal re-encoding of the recalled sequential experience in the cortex. If a remote memory is recalled while other items become active, causal links will be formed between all active items (see Figure 3) in the hippocampus. Hippocampal replays during subsequent offline periods can then alter the cortical representation via the mechanisms described in this paper and thereby mediate reconsolidation.

### Testable predictions

Based on the experiments reported in the Results section, we can make several predictions: (1) If hippocampal projections to cortex are disabled, no cortical replays of recent memories are possible (see Figure 9). (2) The speed of replay of a particular episode increases from one night to the next, even in the absence of additional training, under certain conditions (see Figure 7); namely, the presence of hippocampal learning during SWS and the absence of other experiences during waking. With the rapid passive decay of hippocampal weights, the hippocampal replays tend to become slower with time. If learning is enabled in the hippocampus during SWS, then any replay of the episode would boost the corresponding hippocampal weights. With enough replays (due to the absence of competition from other experiences), the hippocampal weights might become stronger than when the episode was experienced during waking. And as the episode is consolidated in the cortex, the cortex can generate the replays by itself and thereby further boost the strength of the hippocampal replays and vice versa (owing to the bi-directional projections between the hippocampus and the cortex). (3) If there are no other experiences after the exposure to an episode, the decay of the hippocampal memory traces will be significantly slowed as they will be refreshed each night by replays. (4) Recent, salient memories can dominate offline consolidation via replays at the expense of older, less salient memories. Figure 10 shows how an unconsolidated sequence loses the benefit of overnight consolidation due to competition for replays from a more recent experience, causing the hippocampus representation to decay before the cortical representation can develop. Figure 7 illustrates how the more recent sequence comes to dominate the number of replays during sleep.

### Conclusion

In summary, we present a first-of-its-kind computational model of the critical role of the bi-directional interactions between the hippocampus and cortex during wake and sleep for the systems-level consolidation of sequential experiences. This model simulates time-compressed replays of sequences during offline periods that are coordinated between the hippocampus and cortex, and emerge dynamically in response to the recency, frequency, and other saliency factors of the constituent items in the various experiences. Simulation experiments reveal that hippocampal learning during SWS may not be critical for cortical memory consolidation, and provide insights into how salience-driven competition for limited replay opportunities during SWS can underlie the interference among various experiences including those that do not overlap in their content. Further, the model provides an explanation in the context of systems-level consolidation for the neurophysiological evidence that has been argued in favor of the alternate multiple trace theory, as well as for the phenomenon of reconsolidation of remote memories.

## Acknowledgments

This material is based upon work supported by the Defense Advanced Research Projects Agency (DARPA) via the Army Research Office (ARO) under Contract No. W911NF-16-C-0018 (Restoring Active Memory (RAM) Replay) and via Air Force Research Laboratory (AFRL) Contract No. FA8750-18-C-0103 (Lifelong Learning Machines: L2M). Any opinions, findings, and conclusions or recommendations expressed in this material are those of the author(s) and do not necessarily reflect the views of DARPA, ARO, AFRL, or the US Government. We would like to thank Dr. Nicholas Ketz and Dr. Itamar Lerner for their helpful feedback on an earlier version of this manuscript.

